# Birth weight is associated with brain tissue volumes seven decades later, but not with age-associated changes to brain structure

**DOI:** 10.1101/2020.08.27.270033

**Authors:** Emily N. W. Wheater, Susan D. Shenkin, Susana Muñoz Maniega, Maria Valdés Hernández, Joanna M. Wardlaw, Ian J. Deary, Mark E. Bastin, James P. Boardman, Simon R. Cox

## Abstract

Birth weight, an indicator of fetal growth, is associated with cognitive outcomes in early life and risk of metabolic and cardiovascular disease across the life course. Cognitive ability in early life is predictive of cognitive ability in later life. Brain health in older age, defined by MRI features, is associated with cognitive performance. However, little is known about how variation in normal birth weight impacts on brain structure in later life. In a community dwelling cohort of participants in their early seventies we tested the hypothesis that birthweight is associated with the following MRI features: total brain (TB), grey matter (GM) and normal appearing white matter (NAWM) volumes; whiter matter hyperintensity (WMH) volume; a general factor of fractional anisotropy (gFA) and peak width skeletonised mean diffusivity (PSMD) across the white matter skeleton. We also investigated the associations of birthweight with cortical surface area, volume and thickness. Birthweight was positively associated with TB, GM and NAWM volumes in later life (β ≥ 0.194), and with regional cortical surface area but not gFA, PSMD, WMH volume, or cortical volume or thickness. These positive relationships appear to be explained by larger intracranial volume rather than by age-related tissue atrophy, and are independent of body height and weight in adulthood. This suggests that larger birthweight is linked to increased brain tissue reserve in older life, rather than a resilience to age-related changes in brain structure, such as tissue atrophy or WMH volume.

**Significance Statement:** Cognitive brain ageing carries a high personal, societal and financial cost and understanding its developmental origins is important for identifying possible preventative strategies. In a sample of older participants from the Lothian Birth Cohort 1936 we were able to explore the neurobiological correlates of birth weight, which is indicative of the fetal experience. We find that higher birth weight is related to larger brain tissue volumes in later life, but does not modify the trajectory of age-related change. This suggests that early life growth confers preserved differentiation, rather than differential preservation with regards to brain reserve. That these effects are detectable into later life indicates that this variable may be valuable biomarker in the epidemiology of ageing.

## Introduction

The Developmental Origins of Health and Disease hypothesis, formulated by David Barker, posits that the fetal development influences health and disease risk throughout the life course (1). Birth weight, an indicator of fetal growth, is associated with cardiovascular disease risk (2), and neurological and psychiatric outcomes from early life into adulthood (3), as well as early life cognitive (4, 5). Higher cognitive ability in early life is associated with reduced risk of dementia and cognitive decline in later life (6–8). Identifying potential determinants of cognitive and brain aging is of high importance, and the nature of brain reserve that a higher birthweight might confer has been underexplored. It is unclear whether birth weight is a marker of preserved differentiation (differences in how much brain tissue an individual has to begin with), differentiated preservation (differences in the speed of decline), or both (9).

Birth weight within the normal range (2.5 – 4.5 kg) correlates with cognitive performance in childhood (Pearson’s r ~ 0.17) (10), and this influence may extend into adulthood and later life (11, 12). Birth weight also demonstrates a relationship with brain volume (β ~ 0.17), and with a regional patterning of positive associations across the cortical surface area in a cohort of young adults and adolescents - the authors of this study suggested that the regional effects of birth weight on cortical surface area may be due to head size or body size, but this was not explicitly tested (13). Importantly, the association between birth weight and brain tissue volumes and intracranial volume is evident in later life (14, 15). However, the distinct contributions of anthropometric variables such as height, weight and head size remain unclear. Studies on diffusion MRI and birth weight have found white matter alterations in adolescents born with very low birth weight (<1500g), compared to those who were born normal birth weight life (16, 17), and birth weight has been found to have a positive association with frontal white matter fractional anisotropy in later (18). However, little is known about the impact of birth weight variation on white matter microstructure in later life. Below the normal range birth weight is linked to a range of neurobiological correlates that extend into adolescence including reduced white matter microstructural integrity (19), and deviations in cortical thickness (both region dependent thickening and thinning) (20, 21), and volumetric alterations (22) that may be associated with reduced cognitive function (Hadaya and Nosarti, 2020).

To better understand the value of birth weight value as a perinatal indicator of later life brain health, it is important to establish the neurobiological correlates of birth weight into older age. We investigated the following global measures, each of which is reliably associated with cognitive ability in later life with modest effect sizes: total brain (TB) (24, 25), grey matter (GM) and normal appearing white matter (NAWM) which are also linked to cognition (26, 27), as well as white matter hyperintensity (WMH) volume, a marker of cerebrovascular disease in older age, is associated with cognitive decline (26, 28, 29). In older age, brain volume may be affected both by the degree of soft tissue atrophy that has occurred and by maximal healthy brain size (indexed by intracranial volume (ICV), an archaeological measure of maximal brain volume that is invariant with age (25, 30). To investigate the effects of age-related atrophy on associations between birth weight and brain volumes, we controlled for ICV (25). As birth weight is also associated with greater adult height and weight (31), we controlled for these variables to investigate whether a relationship with brain volumes may be due to overall body dimension, rather than to a specific effect on brain or head growth (32). We selected two global measures of white matter microstructure: gFA, which reflects the pattern of covariance of FA values among white matter pathways of the brain and is correlated with preterm birth (33), ageing (34), and cognitive functioning in older age (27), and PSMD, which is effective at capturing diffuse pathology in otherwise normal appearing white matter (36–38). We further investigated regional associations between birth weight and cortical thickness, volume and surface area and explicitly tested the contribution of head size and body size.

To test the hypothesis that birthweight is associated with brain macrostructure and white matter microstructure in older age, we linked these MRI features of brain structure at 73 years of age with perinatal data using a well-characterised single-year-of-birth cohort of healthy community dwelling-older adults (the Lothian Birth Cohort 1936). To understand *how* birth weight has an impact on brain structure we considered the potential roles of i) body size ii) age-related brain tissue atrophy, and iii) cardiovascular and metabolic disease, which is also linked to low birth weight and may mediate the relationship between birth weight and brain structure.

## Results

One hundred and thirty-seven participants of the Lothian Birth Cohort 1936 (LBC1936), a cohort of community dwelling older participants, had both birth weight recorded and structural and diffusion MRI, out of a total of 866 participants in the second wave of data collection (Table 1). In this sample there were no cases of dementia or Parkinson’s disease. Six participants were low birth weight (<2500g); no participants were high birth weight (>4500g). There were no significant differences between those who had both MRI and birth weight data and those who did not in terms of age 11 IQ, age at MRI scan, sex, height, weight, BMI, smoking, hypertension, hypercholesterolemia, diabetes, cardiovascular disease history, or stroke history (all p values >0.1).

**Table 1.** Participant characteristics

|  |  |
| --- | --- |
| Total N | 137 |
| Female/Male | 63/74 |
| Mean age (range) / years | 72.6 (71.1 – 74.1) |
| Mean birth weight (range) /g | 3346 (1843 – 4423) |
| ≤2500 | 6 |
| 2501-3000 | 22 |
| 3001-3500 | 58 |
| 3501-4000 | 40 |
| 4001-4500 | 11 |
| Mean height (range) /cm | 166.0 (146.0 – 185.5) |
| Mean weight (range) / kg | 78.35 (50 – 116.8) |
| Mean BMI (range) | 28.37 (19.32 – 45.65) |
| Smoking (current/ex/never) | 14/55/68 |
| Hypertension (yes/no) | 66/71 |
| Hypercholesterolemia (yes/no) | 62/75 |
| Diabetes diagnosis (yes/no) | 21/116 |
| Cardiovascular disease history (yes/no) | 43/94 |
| Stroke history (yes/no) | 11/126 |

*Note.* Summary of descriptive data for LBC 1936 sample. BMI: body-mass index. Hypertension, hypercholesterolemia, diabetes, cardiovascular and stroke history were obtained by participant self-report in a structured medical interview.

### Birth weight is associated with brain volumes but not atrophy

We first performed linear regression models corrected for age and sex to test the association between birth weight and the following brain MRI features: total brain (TB), grey matter (GM), normal appearing white matter (NAWM), and white matter hyperintensity (WMH) volumes, a general factor of fractional anisotropy (gFA) and peak width skeletonised mean diffusivity (PSMD). The results of age- and sex-corrected associations between birth weight and global brain measures are presented in Table 2. Individuals with a higher birth weight showed generally higher later life TB (β = 0.259, p <0.001), GM (β = 0.194, p = 0.009), and NAWM (β = 0.293, p <0.001) volumes, all of which survived false discovery rate (FDR) multiple comparison correction. However, birth weight was not significantly associated with WMH, or diffusion metrics of white matter microstructure: gFA or PSMD (β ≥ −0.051, p > 0.555).

**Table 2.**
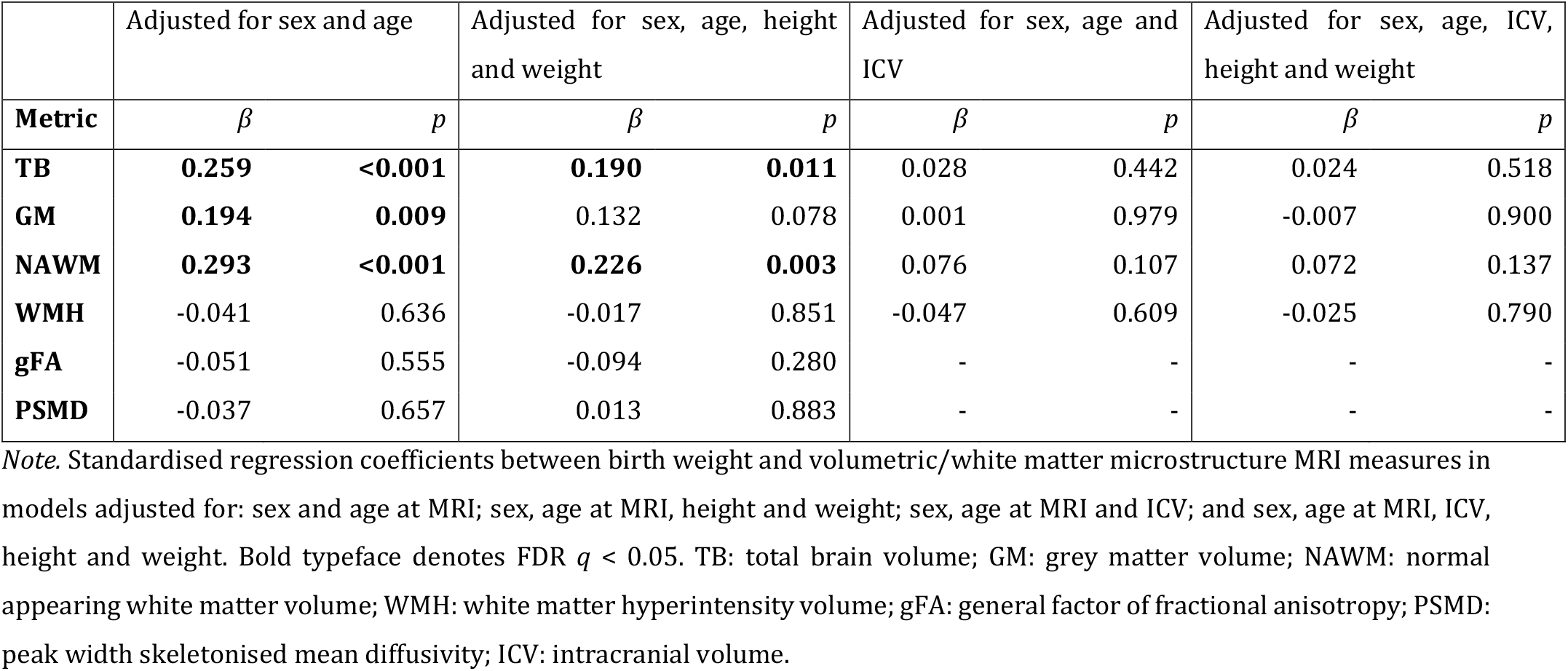
Associations between birth weight and global brain structure at age 73.

|  | Adjusted for sex and age |  | Adjusted for sex, age, height and weight |  | Adjusted for sex, age and ICV |  | Adjusted for sex, age, ICV, height and weight |  |
| --- | --- | --- | --- | --- | --- | --- | --- | --- |
| **Metric** | $\beta$ | $p$ | $\beta$ | $p$ | $\beta$ | $p$ | $\beta$ | $p$ |
| **TB** | **0.259** | **<0.001** | **0.190** | **0.011** | 0.028 | 0.442 | 0.024 | 0.518 |
| **GM** | **0.194** | **0.009** | 0.132 | 0.078 | 0.001 | 0.979 | -0.007 | 0.900 |
| **NAWM** | **0.293** | **<0.001** | **0.226** | **0.003** | 0.076 | 0.107 | 0.072 | 0.137 |
| **WMH** | -0.041 | 0.636 | -0.017 | 0.851 | -0.047 | 0.609 | -0.025 | 0.790 |
| **gFA** | -0.051 | 0.555 | -0.094 | 0.280 | - | - | - | - |
| **PSMD** | -0.037 | 0.657 | 0.013 | 0.883 | - | - | - | - |

*Note.* Standardised regression coefficients between birth weight and volumetric/white matter microstructure MRI measures in models adjusted for: sex and age at MRI; sex, age at MRI, height and weight; sex, age at MRI and ICV; and sex, age at MRI, ICV, height and weight. Bold typeface denotes FDR $q < 0.05$ . TB: total brain volume; GM: grey matter volume; NAWM: normal appearing white matter volume; WMH: white matter hyperintensity volume; gFA: general factor of fractional anisotropy; PSMD: peak width skeletonised mean diffusivity; ICV: intracranial volume.

Next, we investigated whether the effect of birth weight on brain volumes was accounted for by larger body size by controlling for height and weight in addition to age at scan and sex. Following adjustment for height and weight, TB and NAWM volumes remained significantly associated with birth weight, though the strength of the association was reduced by 27% and 23%, respectively. GM volume was no longer significantly associated with birth weight (effect size was attenuated by 31.9%). Birth weight was significantly associated with later life height (Pearson’s r = 0.209, p = 0.006) but not with later life weight (r = 0.137, p = 0.739).

Results for associations between global volumetric measures corrected for ICV are presented in Table 2. ICV is strongly correlated with TB, GM and NAWM volumes (Pearson’s r ≥ 0.835, p <0.001) but not with WMH volume (Pearson’s r = 0.066 p = 0.086; see Supporting Figure S1). When we included ICV as a covariate for all global volumetric measures, we found no significant associations between birth weight and TB volume (β ≤|0.076|, p ≥ 0.107). Results were nearly identical when height and weight were also included in the model (β ≤|0.072|, p ≥ 0.137). The associations magnitudes and p-values were also similar when these global MRI volumes were expressed as a proportion of ICV (Table S1; (β ≤|0.149|, p > 0.070)).

As the inclusion of ICV as a covariate attenuated the relationship between birth weight and brain volumes, we performed an *a posteriori* regression to verify the association between birth weight and ICV directly. ICV was significantly associated with birthweight (β = 0.174; p = 0.009). in a linear regression that included age at MRI, sex, height and weight as covariates.

### Relationship between birth weight and brain structure is independent of cardiovascular risk factors and cardiovascular disease history

When we additionally corrected the associations between birth weight and global brain measures for cardiovascular risk factors and self-report history of cardiovascular disease (Table S2), the initial age- and sex-corrected findings (in Table 2) were modestly attenuated by up to 8.5% (this being the attenuation of effect for NAWM). The associations for birthweight remained significant for TB (β = 0.245, p = 0.00170), GM (β = 0182, p = 0.0188 and NAWM (β = 0.268, p = 0.000754). Associations with WMH, gFA and PSMD remained small and non-significant (β ≤|0.05|, p ≥ 0.5). In a sensitivity analysis including stroke as a covariate did not substantially alter the results (Table S3), though it marginally attenuated the association between GM and birth weight, which was non-significant after FDR correction (B = 0.172, p = 0.031). The size of the attenuation of the effect when stroke was added as a covariate was 5.5%. In a non-stroke sub-group (n = 126) analysis birth weight and GM were still significantly associated (β = 0.203, p = 0.011).

### Associations between birth weight and regional grey matter measures

Vertex wise analysis revealed positive associations between birth weight and surface area on the bilateral temporal (inferior and middle), cingulate (anterior and posterior segments) and anterior frontal (inferior frontal and frontopolar), supramarginal and medial occipital cortices, as well as evidence for associations in the motor and somatosensory cortices and right-sided medial and lateral orbitofrontal, posterior fusiform, angular gyrus and supramarginal gyrus (Figure 1A). These associations were partially attenuated when correcting for height and weight: mean attenuation = 19.21%, SD = 8.67, max = 52.53% (Figure 1B). Controlling for ICV, in a model that was adjusted for age and sex, rendered the regional patterning of birth weight on regional cortical surface area non-significant: mean attenuation = 48.0%, SD = 13.81, max = 98.43% (Figure 1C). There were no significant associations between birth weight and cortical thickness or cortical volume.

**Figure 1.**
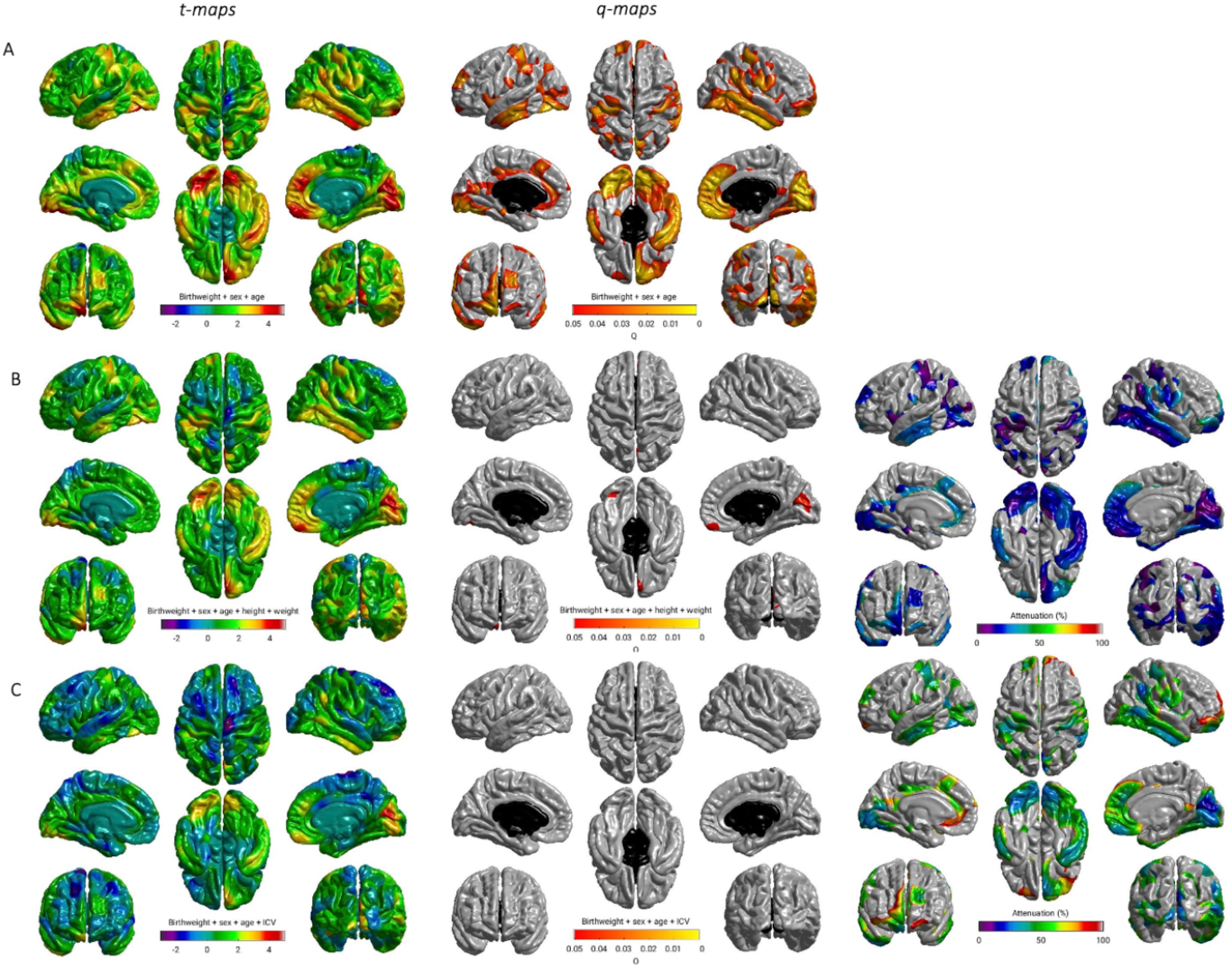
Regional distribution of associations between birth weight and cortical surface area: A) adjusted for sex and age; B) adjusted for age, sex, height and weight; C) adjusted for age, sex, and ICV. T maps (left); FDR q values (middle), far right (B and C) shows the percentage attenuation between the model shown in A, and the additionally adjusted models shown in B and C.

## Discussions

In this well-characterised cohort of older community-dwelling adults, we report that higher birth weight is associated with larger TB, GM and NAWM volumes seven decades later. Importantly, these associations become non-significant with inclusion of ICV as a covariate, whereas associations with WMH, a brain marker of older age neurodegenerative processes among healthy adults, were null. This suggests that the association between birth weight and brain tissue volumes are likely due to the influence of brain size in early life, rather than brain tissue atrophy in later life. This indicates that higher birthweight may confer brain reserve in older age, operating by preserved differentiation rather than differentiated preservation.

Birth weight is positively correlated with adult height and weight (31, 39), and so we controlled for these to test whether the positive associations between birthweight and brain volumes were due to overall body size. Controlling for the effects of these anthropometric indices attenuated the effects between brain volumes and birth weight, but as both TB and NAWM volumes remained significantly positively associated with birth weight when height and weight were controlled for, this relationship appears largely independent of these indices. This was not the case for GM volume, although the effect size attenuation was modest. Given known associations between low birth weight and cardiovascular outcomes (40), and between cardiovascular risk factors and brain structural outcomes (41–43), we included these as covariates in a supplementary analysis and found that, although there was a modest attenuation of the effect (by up to 8.6%), results were robust. In analyses of cortex associations, we found a strikingly similar pattern of positive associations between cortical surface area to that observed, by Walhovd et al. 2012, in a cohort of young adults and adolescents (13). These regional variations of cortical surface area in association with birth weight are in line with the patterning of differential scaling seen as a function of TB volume (44), and may reflect asymmetric tissue scaling with overall brain dimension rather than an altered developmental trajectory (either maladaptive or compensatory) (45). Walhovd et al. 2012 (13) suggested that this effect might be partly driven by being physically larger and here we directly tested this hypothesis. The effect was partially attenuated by controlling for height and weight (by about 20%) which was sufficient for many of the associations across the cortical surface to become non-significant, though several areas still survived FDR correction. The attenuations of effect when controlling for height and weight were modest in comparison to the attenuations observed when controlling for ICV alone. In our analysis of cortical surface area, the magnitude of attenuation was on average around 50% when controlling for ICV. This is consistent, again, with our global brain tissue findings that the associations between birth weight and brain measures 73 years later are attributable to having a bigger head, rather than due to being physically bigger overall. However, like Walhovd et al. 2012 (13), we found no significant relationship between either cortical thickness or volume with birth weight. Taken together, our results indicate a specific contribution of birth weight to maximal healthy brain size that is independent of its relationship to overall body size as measured by height and weight.

Despite consistent findings that low birth weight is associated with diffuse white matter injury from the neonatal period (33) through to early adulthood (46), captured by measures of white matter microstructure, we did not observe a relationship between birth weight and WMH volume or global measures white matter microstructure in late life. However, in our study the birth weight range of the participants was largely normal and an association between birth weight and white matter microstructure was not evident. This supports the theory that the observed associations between low birth weight or preterm birth and white matter microstructure is the product of perturbation during a limited developmental window which coincides with specific biological and environmental exposures, but those susceptibilities are not apparent for individuals in the normal birth weight range (47).

Our study has some limitations. Due to a lack of information on gestational age at birth, we had no way of estimating the prevalence of fetal growth restriction, indicated by being small for gestational age, in our sample. Height and weight were obtained only in older age rather than in midlife for instance, it is possible that head size is simply a more stable indicator of overall body size than height or weight. There are modest decreases in height that occur from midlife onwards and which accelerate around 70 years of age, due to stooping and flattening of intervertebral discs. However, the Pearson’s correlations between birth weight and adult height reported in this sample are of a very similar magnitude to those reported for a large group of men and women in the age group 56-70 years old (48). In this study we have used a relatively small sample due to the availability of detailed birth data. However, our findings with regards to the positive associations between birth weight and brain tissue volumes in later life and regional cortical surface area are consistent with other studies in both adolescents and adults in late life (13–15) We have provided additional analysis of the impact of birth weight on more detailed measures of white matter microstructure and integrity, as well as demonstrating that the regional effects observed in cortical surface area persist into older age and are likely due to the early impact of birth weight on head growth and maximal head size. Within the LBC1936 participants we compared age 11 IQ (as an indicator of potential early life differences) of those with and without birth weight and MRI data, and found no significant difference between the two groups. This was also the case for the covariates included in our analysis, indicating that our sample is representative of the LBC1936 as a whole. This does not preclude the possibility that there were systematic differences in the characteristics of the women who gave birth in hospital in the 1930s compared to those who did not. In addition, the LBC1936 have higher cognitive ability based on their age 11 scores on the Moray House Test relative to the rest of the population that sat the test in 1947 (49, 50). Therefore, generalization of these results to other groups must be performed with caution. We expect that, due to range restriction, the effect sizes reported here are likely to be underestimates of the true effect sizes that would be observed in the wider population. It is, however, striking that these effects relating to variation in birthweight are still detectable among such a relatively healthy group and across seven decades of life, which suggests that this variable may have valuable explanatory power with regards to the epidemiology of cognitive ageing.

Cognitive function and the emergence of cognitive impairment in older age are products of peak cognitive function (preserved differentiation) and the rate of decline that occurs in later life (differential preservation) (9). Our results indicate that higher birth weight contributes to increased brain volumes in later life in a manner consistent with preserved differentiation, rather than differential preservation. This pattern echoes observations of other contributors to brain structure, such as education, and of cognition: cognitive ability in early life is a predictor of cognitive ability in later life, but does not seem to protect against age related cognitive decline or pathology (51), and similarly correlates of brain structure are often found to be predictive of baseline, but not of trajectory of age related decline (52). Maximal healthy brain size, estimated by ICV, has previously been demonstrated to modify the emergence of clinical symptoms for dementia, in a manner consistent with brain volume reserve effects (53), while birth size is significantly correlated with NART performance (an indicator of earlier, crystallized cognitive ability) at 80 years old (11). We found no evidence that the relationship between birth weight and brain volumes was accounted for by brain tissue atrophy, or cardiovascular risk factors in later life. The relationships between birth weight and brain tissue volumes were attenuated by ICV. As the majority of ICV growth occurs in early childhood (achieving 95% of maximal size by 5 years old) we suggest that the relationship is established in early life (54).

Birth weight demonstrates a modest, positive association with brain tissue reserve in later life, which is likely established in early life. It is not a protective factor against either brain tissue atrophy or pathology in later life (indicated by WMH volume and white matter microstructure measured through diffusion MRI). Our results support a developmental origin of brain tissue reserve that survives into old age.

## Materials and Methods

### Participants

Data were collected from the Lothian Birth Cohort 1936 (LBC1936), a longitudinal study of aging comprising individuals who were born in 1936. They mostly took part in the Scottish Mental Survey 1947 (SMS1947) and most were resident in Edinburgh and its surrounding area (the Lothians) at about age 70 years when recruitment for follow-up testing began. The recruitment, brain imaging, and cognitive testing protocols for the LBC1936 have been reported previously in detail (55–57).

At Wave 2, 866 participants returned (mean age = 72.5 years, SD = 0.7 years), 728 of whom underwent brain structural and diffusion MRI, upon which the current study is based. Ethical approval for the LBC1936 study came from the Multi-Centre Research Ethics Committee for Scotland (MREC/01/0/56; 07/MRE00/58) and the Lothian Research Ethics Committee (LREC/2003/2/29). All participants, who were volunteers and received no financial or other reward, completed a written consent form before any testing took place.

### Birth weight

Birth weight was retrieved from original archival hospital records from the time of birth, accessed from the Lothian Health Service Archive at the Centre for Research Collections in the University of Edinburgh. Participants were born at the Edinburgh hospitals Royal Maternity Simpson Memorial Hospital (n = 74), Elsie Inglis Memorial Maternity Hospital (n = 60), Bellshill Maternity Hospital in Lanarkshire (n =2) and Aberdeen Maternity Hospital (n = 1). Birth weight was recorded in the original records in lb and oz and subsequently converted to grams. Analysis was performed with birth weight as a continuous variable.

### Cardiovascular health covariates

During a structured medical history interview at Wave 2, participants were asked whether they had a history of cardiovascular disease or stroke, or had received a diagnosis from a doctor of diabetes, hypercholesterolemia, or hypertension. They were also asked about their smoking history (current/ex/never). BMI (kg/m^2^) was calculated from height (cm) and weight (kg) which were measured at the same time as the medical interview.

### Brain MRI acquisition and processing

Whole-brain structural and diffusion tensor MRI data were acquired by using a 1.5 T GE Signa Horizon scanner (General Electric, Milwaukee, WI, USA) located at the Brain Research Imaging Centre, University of Edinburgh, soon after cognitive testing and plasma collection. T1-, T2-, T2* and FLAIR-weighted MRI sequences were collected and co-registered (voxel size = 1 × 1 × 2 mm). Intracranial (ICV), total brain (TB), grey matter (GM), total white matter and white matter hyperintensity (WMH) volumes were measured using a semi-automated multispectral fusion method for segmentation of brain tissue volumes from the four structural scans, that is, T2-, T1-, T2*- and FLAIR-weighted MRI, (56) (58). Normal-appearing white matter volume (NAWM) was calculated as the difference between total white matter and WMH volumes.

The diffusion MRI protocol employed a single-shot spin-echo echo-planar diffusion-weighted sequence in which diffusion-weighted volumes (b = 1000 s mm^−2^) were acquired in 64 non-collinear directions, together with seven T2-weighted volumes (b = 0 s mm^−2^). This protocol was run with 72 contiguous axial slices with a field of view of 256 × 256 mm, an acquisition matrix of 128 × 128 and 2 mm isotropic voxels (56). From these data we derived a factor of general fractional anisotropy (gFA) from quantitative tractography and peak width skeletonised mean diffusivity (PSMD).

Probabilistic neighbourhood tractography, an automatic tract segmentation method with good reproducibility, was implemented in the TractoR package for R (http://www.tractor-mri.org.uk) (59). The following 12 tracts of interest in each were segmented using this method: the genu and splenium of the corpus callosum, the bilateral rostral cingulum cingulate gyri, the bilateral arcuate, uncinate, and inferior longitudinal fasciculi, and the bilateral anterior thalamic radiations. Tract average FA was derived as the average of all voxels contained within the tract mask weighted by connection probability.

PSMD is the 95^th^ percentile minus the 5^th^ percentile of an individual’s MD within the white matter skeleton. Automatic calculation of PSMD followed the procedure described by Baykara et al. (2016) (36). Diffusion MRI data were processed using the standard Tract-based Spatial Statistics (TBSS) pipeline in FSL. First, all participants’ FA volumes were linearly and non-linearly registered to the standard space FMRIB 1-mm FA template. Second, a white matter skeleton was created from the mean of all registered FA volumes. This was achieved by searching for maximum FA values in directions perpendicular to the local tract direction in the mean FA volume. An FA threshold of 0.2 was applied to the mean FA skeleton to exclude predominantly non-white matter voxels. Third, MD volumes were projected onto the mean FA skeleton and further thresholded at an FA value of 0.3 to reduce CSF partial volume contamination using the skeleton mask provided by Baykara et al. (2016) (36). Finally, PSMD was calculated as the difference between the 95th and 5th percentiles of the voxel-based MD values within each subject’s MD skeleton (37). We opted for this metric instead of a general factor of MD based on prior work indicating its stronger associations with cognitive functioning than gMD (35–37).

Finally, each of the T1-weighted volumes were processed using FreeSurfer v5.1. Following visual quality control in which the outputs for each participant were inspected for aberrant surface meshes, skull stripping and tissue segmentation failures, their estimated cortical surfaces were registered to the ‘fsaverage’ template, yielding a measure of regional volume, surface area and thickness at each of 327,684 vertices across the cortical mantle.

### Statistical Analyses

Unless otherwise stated, all analyses were conducted in R 3.4.3 (R Core Team 2015). We conducted t-tests of continuous variables (age, weight, height, BMI, age 11 IQ) and chi square tests of categorical variables (sex, smoking status, diagnosis of hypertension, hypercholesterolemia or diabetes, cardiovascular disease and stroke history) between the cohort subsample with birth weight data and those without birth weight data. For regression models we report standardised regression coefficients and corrected p-values for multiple comparisons using the False Discovery Rate (FDR; Benjamini and Hochberg, 2007). A general factor of white matter tract FA was calculated using confirmatory factor analysis in *lavaan* (61). A latent factor was estimated from the mean FA of the 12 segmented white matter tracts. Each model included residual correlations between the left and right versions of the bilateral tracts, and also the residual correlation between the splenium and the genu of the corpus callosum. Standardised tract loadings on a general factor of FA (gFA) were > 0.4. Average variance explained = 31.4% Model fit statistics: Tucker Lewis Index= 0.961, Comparative Fit Index = 0.972, root mean square error of approximation = 0.043, standardised root mean square residual = 0.031, X^2^(48)= 106.341, p <0.001. Tract loadings are reported in Table S4. A standardised factor score was extracted from the model for subsequent analysis. WMH volume was log transformed to meet the assumptions of homoscedasticity and normality of error distribution for linear regression.

Initially, we conducted linear regression models of birth weight and global brain image features, with a separate model for each of TB, GM, NAWM and WMH volumes, and PSMD and gFA as the outcome variable. Sex and age in days at MRI acquisition were included as covariates in all models. These models were then conducted with the inclusion of anthropometric covariates height and weight to isolate unique effects of birth weight from late life body size. Pearson correlations were calculated for birth weight with height and weight in later life. Next, because associations between smaller brain volumes in later life with lower birth weight may be explained by increased atrophy in later life, or by having a smaller maximal brain size, prior to age-associated atrophy, we re-ran the associations using the global brain volumes (TB, GM, NAWM, WMH) with ICV as a covariate in addition to age at scan and sex. ICV is a reliable archaeological metric for maximal healthy brain size (25). There are several potential ways of controlling for ICV: one is to express brain volumes as a proportion (i.e. a ratio of volume to ICV), and another is to include ICV as a covariate in analysis (62). For completeness we have included regression results for analysis using the proportion method in supporting information (Table S1).

Due to the relationship between birthweight and cardiovascular disease in later life, and the association between cardiovascular disease and brain structural outcomes, we conducted a set of models adjusting for age and sex and the following health covariates to control for the potential influence on results of cardiovascular and metabolic disease risk: Body Mass Index (BMI), hypertension, diabetes, hypercholesterolemia, smoking status (three categories: current, ex and never), and cardiovascular disease history.

Associations between birth weight and cortical measures (volume, thickness and surface area) were investigated in vertex-wise analyses across cortical mantle, using data smoothed at 20 FWHM to the FreeSurfer average surface (brainstem removed). Age and sex were included as covariates in all models. We additionally included height and weight to control for adult body size. In a separate analysis ICV was included in addition to age and sex to investigate whether the effect of birth weight on brain volumes was accounted for by larger head size/maximal brain size. Cortical volume, thickness and surface area vertex-wise analyses were performed across the average surface with linear models to investigate the effect of birth weight on cortical surface area using the SurfStat toolbox (http://www.math.mcgill.ca/keith/surfstat) for Matrix Laboratory R2018a (The MathWorks Inc., Natick, MA), for which 130 participants had complete MRI, birth weight and covariate data. Results are reported in the form of t-statistics and FDR-corrected q-maps displayed on the ‘fsaverage’ cortical surface.

### Data sharing

LBC 1936 data supporting the findings of this paper are available from the corresponding author upon reasonable request. LBC1936 data are not publicly available due to them containing information that could compromise participant consent and confidentiality. Reasonable requests for original image data will be considered through the Brain Images of Normal Subjects (BRAINS) image bank governance process: www.brainsimagebank.ac.uk (63).

## Supporting information

Wheater(2020)Supporting_Information

## Acknowledgments

ENWW is supported by the Wellcome Trust Translational Neuroscience PhD fellowship programme at the University of Edinburgh (203769/Z/16/A).

The Lothian Birth Cohorts group is funded by Age UK (Disconnected Mind grant) and the Medical Research Council (G0701120, G1001245, MR/M013111/1, MR/R024065/1). SRC, MEB and IJD were also supported by a National Institutes of Health (NIH) research grant R01AG054628

Magnetic Resonance Image acquisition and analyses were conducted at the Brain Research Imaging Centre, Neuroimaging Sciences, University of Edinburgh (www.bric.ed.ac.uk) which is part of SINAPSE (Scottish Imaging Network—A Platform for Scientific Excellence) collaboration (www.sinapse.ac.uk) funded by the Scottish Funding Council and the Chief Scientist Office.

We thank the Lothian Birth Cohort 1936 participants who took part in this study, the Lothian Birth Cohort 1936 research team members, and radiographers at the Brain Research Imaging Centre

## References

1. D. J. P. Barker, The Developmental Origins of Adult Disease. J. Am. Coll. Nutr. 23, 588S–595S (2004).

2. J. J. M. Geelhoed, V. W. V. Jaddoe, Early influences on cardiovascular and renal development. Eur. J. Epidemiol. 25, 677–692 (2010).

3. K. J. O’Donnell, M. J. Meaney, Fetal origins of mental health: The developmental origins of health and disease hypothesis. Am. J. Psychiatry 174, 319–328 (2017).

4. C. Sacchi, et al., Association of Intrauterine Growth Restriction and Small for Gestational Age Status with Childhood Cognitive Outcomes: A Systematic Review and Meta-analysis. JAMA Pediatr. 174, 772–781 (2020).

5. S. D. Shenkin, et al., Birth weight and cognitive function at age 11 years: The Scottish Mental Survey 1932. Arch. Dis. Child. 85, 189–195 (2001).

6. S. Dekhtyar, et al., A life-course study of cognitive reserve in dementia - From childhood to old age. Am. J. Geriatr. Psychiatry 23, 885–896 (2015).

7. V. J. Bourne, H. C. Fox, I. J. Deary, L. J. Whalley, Does childhood intelligence predict variation in cognitive change in later life? Pers. Individ. Dif. 42, 1551–1559 (2007).

8. M. Richards, B. Shipley, R. Fuhrer, M. E. J. Wadsworth, Cognitive ability in childhood and cognitive decline in mid-life: Longitudinal birth cohort study. Br. Med. J. 328, 552–554 (2004).

9. E. M. Tucker-Drob, Cognitive Aging and Dementia: A Life-Span Perspective. Annu. Rev. Dev. Psychol. 1, 177–196 (2019).

10. S. D. Shenkin, J. M. Starr, I. J. Deary, Birth weight and cognitive ability in childhood: A systematic review. Psychol. Bull. 130, 989–1013 (2004).

11. S. D. Shenkin, I. J. Deary, J. M. Starr, Birth parameters and cognitive ability in older age: a follow-up study of people born 1921-1926. Gerontology 55, 92–98 (2009).

12. B. J. Grove, S. J. Lim, C. R. Gale, S. D. Shenkin, Birth weight and cognitive ability in adulthood: A systematic review and meta-analysis. Intelligence 61, 146–158 (2017).

13. K. B. Walhovd, et al., Long-term influence of normal variation in neonatal characteristics on human brain development. Proc. Natl. Acad. Sci. U. S. A. 109, 20089–20094 (2012).

14. M. Muller, et al., Late-life brain volume: a life-course approach. The AGES-Reykjavik study. Neurobiol. Aging 41, 86–92 (2016).

15. M. Muller, et al., Birth size and brain function 75 years later. Pediatrics 134, 761–770 (2014).

16. J. Skranes, et al., Clinical findings and white matter abnormalities seen on diffusion tensor imaging in adolescents with very low birth weight. Brain 130, 654–666 (2007).

17. T. R. Vangberg, et al., Changes in white matter diffusion anisotropy in adolescents born prematurely. Neuroimage 32, 1538–1548 (2006).

18. S. D. Shenkin, et al., Birth parameters are associated with late-life white matter integrity in community-dwelling older people. Stroke 40, 1225–1228 (2009).

19. L. Eikenes, G. C. Løhaugen, A. M. Brubakk, J. Skranes, A. K. Håberg, Young adults born preterm with very low birth weight demonstrate widespread white matter alterations on brain DTI. Neuroimage 54, 1774–1785 (2011).

20. H. M. A. de Bie, et al., Global and regional differences in brain anatomy of young children born small for gestational age. PLoS One 6 (2011).

21. K. J. Bjuland, G. C. C. Løhaugen, M. Martinussen, J. Skranes, Cortical thickness and cognition in very-low-birth-weight late teenagers. Early Hum. Dev. 89, 371–380 (2013).

22. V. R. Karolis, et al., Volumetric grey matter alterations in adolescents and adults born very preterm suggest accelerated brain maturation. Neuroimage 163, 379–389 (2017).

23. L. Hadaya, C. Nosarti, The neurobiological correlates of cognitive outcomes in adolescence and adulthood following very preterm birth. Semin. Fetal Neonatal Med. 25, 101117 (2020).

24. J. Pietschnig, L. Penke, J. M. Wicherts, M. Zeiler, M. Voracek, Meta-analysis of associations between human brain volume and intelligence differences: How strong are they and what do they mean? Neurosci. Biobehav. Rev. 57, 411–432 (2015).

25. N. A. Royle, et al., Estimated maximal and current brain volume predict cognitive ability in old age. Neurobiol. Aging 34, 2726–2733 (2013).

26. Z. Arvanitakis, et al., Association of white matter hyperintensities and gray matter volume with cognition in older individuals without cognitive impairment. Brain Struct. Funct. 221, 2135–2146 (2016).

27. S. R. Cox, C. Fawns-Ritchie, E. M. Tucker-Drob, I. J. Deary, Brain imaging correlates of general intelligence in UK Biobank. Intelligence 76, 101376 (2019).

28. R. Au, et al., Association of white matter hyperintensity volume with decreased cognitive functioning: The Framingham Heart Study. Arch. Neurol. 63, 246–250 (2006).

29. S. J. Wiseman, et al., Cognitive abilities, brain white matter hyperintensity volume, and structural network connectivity in older age. Hum. Brain Mapp. 39, 622–632 (2018).

30. D. D. Blatter, et al., Quantitative volumetric analysis of brain MR: Normative database spanning 5 decades of life. Am. J. Neuroradiol. 16, 241–251 (1995).

31. H. T. Sørensen, et al., Birth weight and length as predictors for adult height. Am. J. Epidemiol. 149, 726–729 (1999).

32. L. M. O’Brien, et al., Adjustment for whole brain and cranial size in volumetric brain studies: A review of common adjustment factors and statistical methods. Harv. Rev. Psychiatry 14, 141–151 (2006).

33. E. J. Telford, et al., A latent measure explains substantial variance in white matter microstructure across the newborn human brain. Brain Struct. Funct. 222 (2017).

34. S. R. Cox, et al., Ageing and brain white matter structure in 3,513 UK Biobank participants. Nat. Publ. Gr. 7, 13629 (2016).

35. L. Penke, et al., A General Factor of Brain White Matter Integrity Predicts Information Processing Speed in Healthy Older People. J. Neurosci. 30, 7569–7574 (2010).

36. E. Baykara, et al., A Novel Imaging Marker for Small Vessel Disease Based on Skeletonization of White Matter Tracts and Diffusion Histograms. Ann. Neurol. 80, 581–592 (2016).

37. I. J. Deary, et al., Brain Peak Width of Skeletonized Mean Diffusivity (PSMD) and Cognitive Function in Later Life. Front. Psychiatry 10, 1–10 (2019).

38. C. Vinciguerra, et al., Peak width of skeletonized mean diffusivity (PSMD) as marker of widespread white matter tissue damage in multiple sclerosis. Mult. Scler. Relat. Disord. 27, 294–297 (2019).

39. M. G. Eide, et al., Size at birth and gestational age as predictors of adult height and weight. Epidemiology 16, 175–181 (2005).

40. C. Osmond, D. J. P. Barker, P. D. Winter, C. H. D. Fall, S. J. Simmonds, Early growth and death from cardiovascular disease in women. Br. Med. J. 307, 1519–1524 (1993).

41. S. R. Cox, et al., Associations between vascular risk factors and brain MRI indices in UK Biobank. Eur. Heart J. 40, 2290–2299 (2019).

42. S. Debette, et al., Midlife vascular risk factor exposure accelerates structural brain aging and cognitive decline. Neurology 77, 461–468 (2011).

43. R. Song, et al., Associations Between Cardiovascular Risk, Structural Brain Changes, and Cognitive Decline. J. Am. Coll. Cardiol. 75, 2525–2534 (2020).

44. P. K. Reardon, et al., Normative brain size variation and brain shape diversity in humans. Science (80-.). 360, 1222–1227 (2018).

45. L. Jäncke, F. Liem, S. Merillat, Weak correlations between body height and several brain metrics in healthy elderly subjects. Eur. J. Neurosci. 50, 3578–3589 (2019).

46. M. J. Pascoe, T. R. Melzer, L. J. Horwood, L. J. Woodward, B. A. Darlow, Altered grey matter volume, perfusion and white matter integrity in very low birthweight adults. NeuroImage Clin. 22, 101780 (2019).

47. J. P. Boardman, S. J. Counsell, Invited Review: Factors associated with atypical brain development in preterm infants: insights from magnetic resonance imaging. Neuropathol. Appl. Neurobiol. 46, 413–421 (2020).

48. H. Ylihärsilä, et al., Birth size, adult body composition and muscle strength in later life. Int. J. Obes. 31, 1392–1399 (2007).

49. W. Johnson, I. J. Deary, Placing inspection time, reaction time, and perceptual speed in the broader context of cognitive ability: The VPR model in the Lothian Birth Cohort 1936. Intelligence 39, 405–417 (2011).

50. W. Johnson, A. J. Gow, J. Corley, J. M. Starr, I. J. Deary, Location in cognitive and residential space at age 70 reflects a lifelong trait over parental and environmental circumstances: The Lothian Birth Cohort 1936. Intelligence 38, 402–411 (2010).

51. A. J. Gow, et al., Stability and Change in Intelligence From Age 11 to Ages 70, 79, and 87: The Lothian Birth Cohorts of 1921 and 1936. Psychol. Aging 26, 232–240 (2011).

52. S. J. Ritchie, et al., Risk and protective factors for structural brain ageing in the eighth decade of life. Brain Struct. Funct. 222, 3477–3490 (2017).

53. H. Wolf, P. Julin, H. J. Gertz, B. Winblad, L. O. Wahlund, Intracranial volume in mild cognitive impairment, Alzheimer’s disease and vascular dementia: Evidence for brain reserve? Int. J. Geriatr. Psychiatry 19, 995–1007 (2004).

54. S. Sgouros, A. D. Hockley, J. H. Goldin, M. J. C. Wake, K. Natarajan, Intracranial volume change in craniosynostosis. J. Neurosurg. 91, 617–625 (1999).

55. I. J. Deary, et al., The Lothian Birth Cohort 1936: a study to examine influences on cognitive ageing from age 11 to age 70 and beyond. BMC Geriatr. 12, 1–12 (2007).

56. J. M. Wardlaw, et al., Brain aging, cognition in youth and old age and vascular disease in the Lothian Birth Cohort 1936: Rationale, design and methodology of the imaging protocol. Int. J. Stroke 6, 547–559 (2011).

57. A. M. Taylor, A. Pattie, I. J. Deary, Cohort Profile Update: The Lothian Birth Cohorts of 1921 and 1936. Int. J. Epidemiol. 47 (2018).

58. M. D. C. Valdés Hernández, K. J. Ferguson, F. M. Chappell, J. M. Wardlaw, New multispectral MRI data fusion technique for white matter lesion segmentation: Method and comparison with thresholding in FLAIR images. Eur. Radiol. 20, 1684–1691 (2010).

59. J. D. Clayden, et al., TractoR: Magnetic Resonance Imaging and Tractography with R. J. Stat. Softw. 44, 1–18 (2011).

60. Y. Benjamini, Y. Hochberg, Controlling the False Discovery Rate: A Practical and Powerful Approach to Multiple Testing. J. R. Stat. Soc. Ser. B (Statistical Methodol. 57, 289–300 (2007).

61. Y. Rosseel, lavaan: An R Package fot Structural Equation Modeling. J. Stat. Softw. 48 (2012).

62. L. M. O’Brien, et al., Statistical adjustments for brain size in volumetric neuroimaging studies: Some practical implications in methods. Psychiatry Res. 193, 113–122 (2011).

63. D. E. Job, et al., A brain imaging repository of normal structural MRI across the life course: Brain Images of Normal Subjects (BRAINS). Neuroimage 144, 299–304 (2017).

