## Supplementary material for "Birth weight is associated with brain tissue volumes seven decades later, but not with age-associated changes to brain structure": Wheater(2020)Supporting_Information

**This PDF file includes:**

Tables S1 to S4
Figure S1

**Table S1.** Associations between birth weight and the ratio of global brain volumetric MRI measures as a proportion of intracranial volume, correcting for age and sex

| | $\beta$ | $p$ |
|---|---|---|
| TB/icv | 0.006 | 0.937 |
| GM/icv | -0.074 | 0.379 |
| NAWM/icv | 0.149 | 0.080 |
| WMH/icv | -0.094 | 0.273 |

*Note.* Standardised regression coefficients between birth weight and volumetric MRI measures expressed as a ratio with ICV. Bold typeface denotes FDR $q < 0.05$. TB: total brain volume; GM: grey matter volume; NAWM: normal appearing white matter volume; WMH: white matter hyperintensity volume.

**Table S2.** Associations between birth weight and volumetric MRI measures correcting for age, sex, cardiovascular risk factors and cardiovascular disease history.

| | $\beta$ | $p$ |
|---|---:|---:|
| TB | **0.245** | **0.002** |
| GM | **0.182** | **0.019** |
| NAWM | **0.268** | **<0.001** |
| WMH | -0.035 | 0.701 |
| gFA | -0.053 | 0.547 |
| PSMD | -0.016 | 0.850 |

*Note.* Standardised regression coefficients between birth weight and volumetric/white matter microstructure MRI measures. Bold typeface denotes FDR $q < 0.05$. TB: total brain volume; GM: grey matter volume; NAWM: normal appearing white matter volume; WMH: white matter hyperintensity volume; gFA: general factor of fractional anisotropy; PSMD: peak width skeletonised mean diffusivity.

**Table S3.** Associations between birth weight and volumetric MRI measures correcting for age, sex, cardiovascular risk factors and cardiovascular disease and stroke history.

| | $\beta$ | $p$ |
|---|---:|---:|
| TB | **0.233** | **0.004** |
| GM | 0.172 | 0.031 |
| NAWM | **0.262** | **0.001** |
| WMH | -0.049 | 0.598 |
| gFA | -0.060 | 0.503 |
| PSMD | 0.010 | 0.910 |

*Note.* Standardised regression coefficients between birth weight and volumetric/white matter microstructure MRI measures. Bold typeface denotes FDR $q < 0.05$. TB: total brain volume; GM: grey matter volume; NAWM: normal appearing white matter volume; WMH: white matter hyperintensity volume; gFA: general factor of fractional anisotropy; PSMD: peak width skeletonised mean diffusivity.

**Table S4.** Tract loadings on general factor of fractional anisotropy

| Tract | Standardised loadings. |
|---:|---:|
| Splenium | 0.425 |
| Genu | 0.624 |
| LArc | 0.613 |
| RArc | 0.606 |
| LATR | 0.600 |
| RATR | 0.585 |
| LCing | 0.520 |
| RCing | 0.530 |
| LUnc | 0.634 |
| RUnc | 0.661 |
| LILF | 0.437 |
| RILF | 0.416 |

*Note.* the genu and splenium of the corpus callosum, LArc – Left arcuate, RArc – right arcuate, LATR – left anterior thalamic radiation, RATR – right anterior thalamic radiation, LCing – Left cingulum bundle, RCing – Right cingulum bundle, LUnc – left uncinate, RUnc – right uncinate, LILF – left inferior longitudinal fasciculus, RILF – right inferior longitudinal fasciculus.

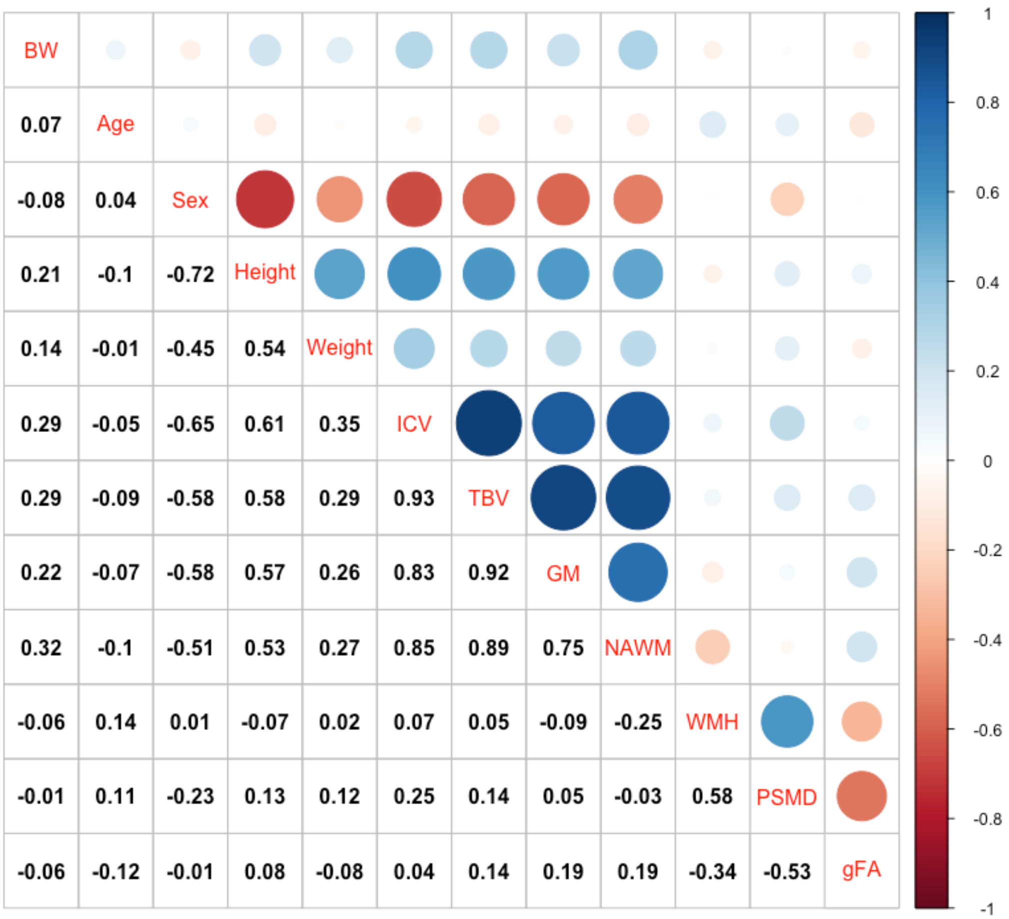


**Figure S1.** Correlation matrix showing Pearson's r correlations between brain MRI features and birth weight, age at MRI, sex, height and weight.

Note BW: birth weight; ICV: intracranial volume; TBV: total brain volume; GM: grey matter volume; NAWM: normal appearing white matter; WMH: white matter hyperintensity volume; PSMD: peak width skeletonized mean diffusivity; gFA: general factor of fractional anisotropy.
